# Increased number and activity of a lateral subpopulation of hypothalamic orexin/hypocretin neurons underlies the expression of an addicted state in rats

**DOI:** 10.1101/356220

**Authors:** Morgan H James, Colin M Stopper, Benjamin A Zimmer, Nikki E Koll, Hannah E Bowrey, Gary Aston-Jones

**Affiliations:** Brain Health Institute, Rutgers University and Rutgers Biomedical and Health Sciences, Piscataway, New Jersey.; The Florey Institute of Neuroscience and Mental Health, Parkville, VIC 3052, Australia; Save Sight Institute, The University of Sydney, Sydney, NSW, 2000, Australia

**Keywords:** Cocaine, plasticity, hypothalamus, behavioral economics, intermittent access, long access

## Abstract

**Background:** The orexin system is important for reward-driven motivation but has not been implicated in the expression of a multi-phenotype addicted state.

**Methods:** Rats were assessed for economic demand for cocaine prior to and following 14d of short- (ShA), long- (LgA) or intermittent-access (IntA) to cocaine. Rats were also assessed for a number of other DSM-V-relevant addiction criteria following differential access conditions. Orexin system function was assessed by i) quantification of numbers and activity of orexin cells, ii) pharmacological blockade of the orexin-1 receptor, and iii) subregion-specific knockdown of orexin cell populations.

**Results:** IntA produced a cluster of addiction-like behaviors that closely recapitulate key diagnostic criteria for addiction to a greater extent than LgA or ShA. IntA was associated with plasticity in orexin cell function, including increased number and activity of orexin-expressing neurons within the lateral hypothalamic (LH) subregion. This plasticity persisted during protracted withdrawal from cocaine for at least 6 months and was associated with enhanced incubation of craving. Selective knockdown of LH orexin neurons reversed the addicted state, and orexin-1 receptor signaling played a larger role in drug seeking after IntA.

**Conclusions:** These data provide the first evidence that LH orexin system function extends beyond general reward seeking to play a critical role in the expression of a multi-phenotype addicted-like state. Thus, the orexin/hypocretin system is a potential novel target for pharmacotherapies designed to treat cocaine addiction. In addition, these data point to the IntA model as a preferred approach to modeling addiction-like behavior in rats.

## Introduction

The hypothalamic orexin (hypocretin) system is a promising target for pharmacotherapies to treat addiction (1–5). Orexin peptides are produced in neurons in caudal hypothalamus, including dorsomedial (DMH), perifornical (PF) and lateral hypothalamus (LH) proper (6, 7); evidence indicates that LH orexin cells in particular underlie drug-seeking behaviors. Indeed, LH orexin neurons are preferentially activated by cocaine- or morphine-associated stimuli, and stimulation of LH orexin neurons drives drug seeking (4, 8–10). This is in contrast to DMH/PF orexin neurons that are activated by stress, waking or general arousal (11–13).

Pharmacological studies support a role for the orexin system in drug seeking, as compounds that block signaling at the orexin-1 receptor (OxR1) reduce seeking of multiple drugs of abuse (1). Moreover, orexin peptides regulate cocaine-induced plasticity at midbrain dopamine synapses (14), and alcohol or cocaine can change orexin peptide or receptor mRNA levels, respectively (15, 16). Importantly however, these behavioral and plasticity studies have almost exclusively utilized limited access models of drug self-administration that do not recapitulate pathological drug seeking/taking in human addicts. Further, these studies typically focus on single addiction endophenotypes, such as sensitization or reinstatement, which do not manifest in isolation in addicts (17). Thus, a role for orexin in a clinically relevant, multifaceted addicted state characterized by disordered and pathological drug seeking has not been reported.

The transition from casual drug-use to an addicted state in humans is often attributed to a gradual escalation of drug intake (tolerance, or shift in hedonic set point) with repeated use (18). Thus, to promote an addiction-like phenotype, laboratory animals are often given unrestricted, continuous access to cocaine in extended sessions (≥6h; long access, LgA) (19). Recent evidence however, indicates that the *pattern* of drug use, rather than the *amount* of drug intake, might be more important for addiction in humans. Experienced cocaine users report an intermittent pattern of intake, where even within a single binge cocaine uses are separated by significant periods of time, resulting in a ‘spiking’ pattern of brain-cocaine levels (20, 21). This spiking pattern can be mimicked in rats by giving intermittent access (IntA) to cocaine in daily sessions; this pattern of intake was associated with enhanced drug motivation (22). Similarly, incentive sensitization develops with prolonged IntA to cocaine (23). It is unclear however, how the IntA model compares to the prototypical LgA model in terms of promoting DSM-V behaviors diagnostic for substance abuse, including increased drug motivation, escalation of intake, continued use despite negative consequences (compulsive intake), and development of dependence.

Here, we used a behavioral economics task to measure changes in demand and ‘hedonic set point’ following IntA versus LgA or ShA. We hypothesized that the well-documented shift in hedonic set point following LgA (19) does not necessarily confer compulsive intake, enhanced reinstatement of responding, or persistently increased motivation for cocaine. In contrast, we predicted that IntA would be superior to the LgA model for promoting these and other elements of a persistent and comprehensive addicted-like state that reflects key clinical features of the human condition. We further hypothesized that this IntA-induced addicted-like state would be accompanied by persistent plasticity of orexin neurons, particularly those in the LH subpopulation, and that the addiction phenotype could be reversed by selective knockdown of these cells or pharmacological blockade of OxR1 signaling.

## Methods and Materials

All experiments were conducted in accordance with procedures established by the Institutional Animal Care and Use Committee of Rutgers University and the *Guide for the care and use of laboratory animals* (National Research Council, 2011).

### Intermittent, long and short access paradigms

In a single IntA session, cocaine was available in twelve 5-min epochs that were separated by 25min periods during which the levers were retracted. During the epochs of cocaine availability, responding was reinforced on an FR1 schedule, whereby each response resulted in a 1s infusion of drug (0.055mg) which was paired with simultaneous (1s) light+tone cues. As per previous reports (22), no timeout periods were imposed during the 5 min epochs when cocaine was available, apart from the time during which the pump was active. To signal the return of cocaine availability following each 25min period of cocaine non availability, the light+tone cues were presented for 5s and animals were given a single priming infusion of cocaine (1s,0.055mg;iv), before the levers were reinserted into the operant chamber. IntA continued for a total of 14d. Rats in the LgA group were given continuous access to cocaine (0.2mg) on an FR1 schedule during daily 6h sessions for 14ds. ShA consisted of continuous access to cocaine (0.2mg) on an FR1 schedule during 1h daily sessions for 14d. In both the ShA and LgA paradigms, cocaine infusions (3.6s) were paired with simultaneous light+tone cues, followed by a 20s timeout period signaled by extinguishing the houselight. Animals were trained on the IntA/LgA/ShA paradigms 5-6d/week.

### Behavioral economics (BE) procedure

To assess economic demand for cocaine, rats were trained on a within-session BE threshold procedure (24) previously described by our laboratory (25–27). In 110min sessions, rats received access to decreasing doses of cocaine in successive 10min intervals on a quarter logarithmic scale (383.5, 215.6, 121.3, 68.2, 38.3, 21.6, 12.1, 6.8, 3.8, 2.2 and 1.2 μg/infusion), which was achieved by decreasing the duration of the pump infusion. For the duration of each infusion, the houselight was turned off and light+tone cues were presented. During this time, presses on the active lever were recorded but did not elicit a second infusion, and a new infusion could be initiated as soon as the previous infusion was completed. By fitting an exponential demand equation to the data(28), we determined a (an inverse measure of motivation), calculated Pmax (the maximum effort the animal is willing to exert to defend preferred brain-cocaine levels), and Q_0_ (consumption at null cost) values (see ‘Demand curve fitting’, *Supplemental Methods and Materials*).

### Test of compulsive responding

Compulsive responding for drug was assessed using a punished responding procedure as previously described (27). A relatively high cocaine infusion dose (0.38mg;2.6sec) was held constant throughout the session; shock was omitted for the first two 10min bins, however, beginning in the third bin, footshocks were delivered during each cocaine infusion and these increased in amplitude every 10min on a tenth- logio scale: 0.13, 0.16, 0.20, 0.25, 0.32, 0.40, 0.50, 0.63, and 0.79 milliamps. Shock resistance was characterized as the maximum cumulative charge in millicoulombs (mC) an animal self-administered in any one bin.

### Morpholino antisense injections

Injector cannulae (28G;Plastics One) were lowered through and 2mm below guide cannulae directed at either LH or PF/DMH; animals received a microinfusion of orexin morpholino antisense (OX-AS) (0.15nmol/0.3μl in 0.5mM phosphate buffer, Gene Tools, 5’-GTATCTTCGGTGCAGTGGTCCAAAT- 3’) or control antisense with the reverse sequence to the OX-AS. These compounds have previously been shown not to affect the expression of non-orexin peptides in nearby neurons(8, 29).

### Experimental overview

Following self-administration training, rats were run on the threshold procedure daily for a minimum of 6d and until the last three sessions produced stable a and Q_0_ values (<25% variability); these values were used as baseline values. Rats were then trained on IntA/LgA/ShA and then immediately re-tested on the threshold procedure. Tests with the OxR1 antagonist SB-334867 (SB) began once animals achieved stability, and there was a minimum of 3 sessions between tests (0,10,30mg/kg, i.p.). Rats were then tested for compulsive responding. In a subset of animals that maintained catheter patency, values from a threshold BE test 50-55d following the final IntA/LgA/ShA session (‘d50’) were also obtained; these animals were then treated with vehicle and SB30 (these rats were not used for any further tests). Following the compulsive responding test, rats underwent extinction training and were tested for cued and/or primed reinstatement under SB conditions (0,10,30mg/kg). A subgroup of rats then underwent 4w of homecage abstinence, at which time they were tested for anxiety- and depression-like behavior. Following 3m abstinence, incubation of craving was assessed under cued reinstatement conditions. Another subgroup of rats were tested for locomotor activity following SB (0,10,30mg/kg). For Fos and orexin cell count studies, rats were returned to the self-administration room (but remained in their homecages) either 1d or 150d following IntA/ShA training and perfused 90mins later. For morpholino studies, rats were tested for cocaine demand each day following antisense infusions; demand values presented are those collected 6d after infusion, as this is the time of peak orexin knockdown (8, 29). Details of animals, drug preparations, surgical procedures, extinction and reinstatement testing, mood assays, histology, imaging and cell counts, locomotor testing, and statistical analyses are included in *Supplemental Methods and Materials.*

## Results

### IntA to cocaine promotes a multi-phenotype addicted state

Escalation of first-hour drug consumption was observed in IntA (F_13,335_ = 6.79, p<0.0001) and LgA (F_13,337_ = 2.9 2 2, p=0.0115) animals, but not ShA animals (p>0.05;**Figure 1a**). Overall escalation of first-hour drug intake was similar between IntA and LgA animals (**Figure 1b and Figure S1a-b**). IntA was associated with an increase in non-rewarded responding across the training period that was not observed following LgA or ShA (p’s<0.05;**Figure S1d-e**). This stronger drug-seeking profile in IntA animals was despite much higher total cocaine consumption in LgA animals (F_2,78_=200.3,p<0.0001;**Figure S1f**).

**Figure 1:**
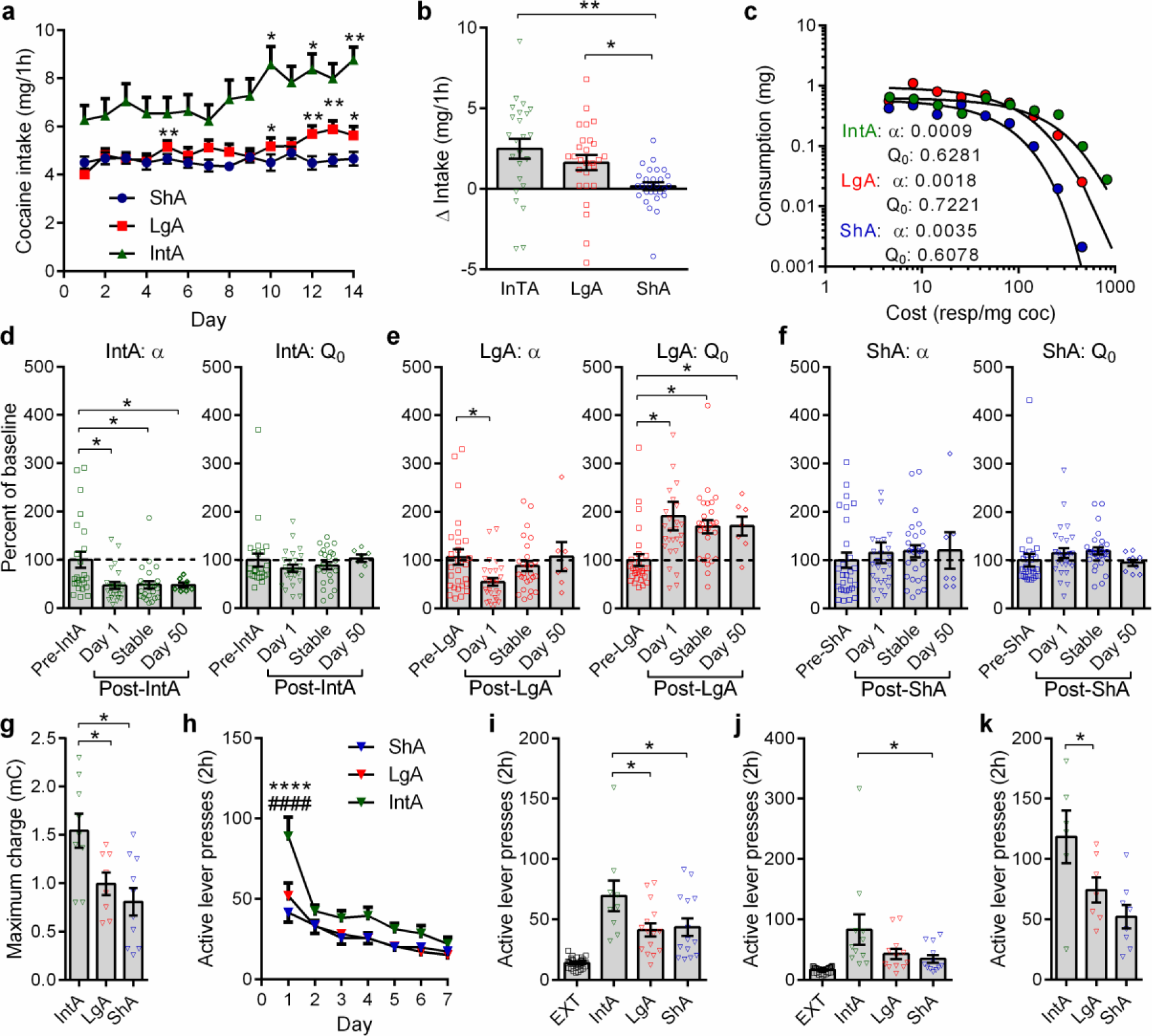
IntA produces a comprehensive addiction-like state. (**a**) IntA animals escalated their intake of cocaine across training, F_13,335_=6.788, p<0.0001. LgA animals also exhibited first-hour escalation of intake, F_13,377_= 2.9 2 2, p=0.0115. There was no change in cocaine intake in ShA rats, F_13_,_391_=0.7945, P=0.5567. Stars denote significant difference relative to Day 1, Holm-Sidak multiple comparisons. IntA: n=24; LgA: n=27; ShA: n=28. (**b**) Delta intake (dl4 minus dl) was similar between IntA and LgA rats and was significantly greater than ShA rats in both cases, F_2,76_=6.805, p=.00l9, Holm-Sidak post-hoc tests. IntA: n= 24; LgA: n=27;ShA:n=28. (**c**) Representative individual demand curves for treatment groups. (**d**) IntA was associated with decreased a values (increased motivation) at ld (t=2.626, p=0.0298; Holm-Sidak post-hoc test), ~7d (stable; t=2.445, p=.0298; Holm-Sidak post-hoc test), and 50d (Wilcoxon test=-24, p=0.0469) following IntA. No changes were observed in Q_0_ at any timepoint. Dl, stable: n=24; D50: n=7. (**e**) LgA only transiently decreased a (Dl; t=2.434, p=0.0438; Holm-Sidak post-hoc test), despite the hedonic set point for cocaine (Q_0_) being increased for up to 50d (t=0.3598, p=.0499; Holm-Sidak post-hoc test). Dl, stable: n=27; D50: n=7. (**f**) ShA had no effect on a or Q_0_ values at any of the timepoints measured. Dl, stable: n=28; D50: n=7. (**g**) IntA rats accepted a higher maximum electrical charge to defend their preferred brain-cocaine levels. mC: millicoulombs (versus LgA, t=2.520, p=0.0l88; versus ShA, t=3.563, p=0.0032; Holm- Sidak post-hoc comparisons). IntA: n=9; LgA: n=8; ShA: n=l0. (**h**) Active lever responding on the first day of extinction training was highest in IntA rats, p<.000l compared to both LgA and ShA rats, Holm-Sidak post-hoc comparisons. IntA: n=26; LgA: n=23; ShA: n=20. (**i**) Cued-reinstatement following extinction training was significantly higher in IntA animals compared to LgA (t=2.398, p=0.0434) and ShA groups (t=2.l78, p=0.0434; Holm-Sidak post-hoc tests). EXT: pooled extinction values across all groups. IntA; n=9; LgA: n=l5; ShA: n=l4. (**j**) Cocaine primed-reinstatement following extinction training was highest in IntA animals (versus ShA, Dunn’s rank 10.84, p=0.0224, post-hoc test). EXT: pooled extinction values across all groups. IntA:n=9;LgA: n=15; ShA:n=14. (**k**) Cued-reinstatement following 12 weeks of home cage abstinence was highest in IntA rats (versus ShA, t=2.168, p=0.0438, Holm-Sidak post-hoc test). IntA: n=6; LgA: n=7; ShA: n=8. Error bars are s.e.m. *P<0.05. **P<0.01 ***P<0.001. ****P<0.0001 vs ShA. ####P<0.0001 vs LgA.

To assess motivation for drug, we tested animals on a behavioral economics task that measures consumption of cocaine across increasing prices (23, 26, 27). Baseline β (an inverse index of motivation) and Q_0_ (cocaine consumption at null cost) values were similar across groups prior to ShA, LgA or IntA (p’s>0.05;**Figure S1g-h**). IntA was associated with a robust decrease in β (increased motivation; F_2,71_=6.275,p=0.0l76;**Figure 1c,d**) that persisted for at least 50d (Wilcoxon test, W=-24,p=0.0469), but no change in Q_0_ (p’s>0.05;**Figure 1d**). In contrast, LgA was associated with only a transient decrease in β (F_2,80_=5.050,p=0.0158;**Figure 1e**) that dissipated within lwk, despite a persistent increase in Q (p’s<0.05;**Figure 1e**). This indicates that tolerance to the hedonic effects of cocaine (reflected here in increased Q_0_, and thought to underlie escalation of intake following LgA) does not confer persistently increased motivation for drug, as has previously been suggested (l9). No changes in a or Q_0_ were observed in ShA animals (**Figure 1f**).

In a test of punished responding for drug, IntA rats accepted a higher maximum electrical charge to defend their preferred brain-cocaine levels compared to LgA or ShA subjects (F_2,24_=6.717,p=0.0048;**Figure 1g**), revealing the development of a strong compulsive drug-seeking phenotype in IntA animals.

IntA rats exhibited higher levels of drug seeking on the first day of extinction training (F_12,462_= 2.643, p=0.0020;**Figure 1h**) and on tests of cue-(F_2,37_=3.270,p=0.0499;**Figure 1i**) or cocaine-primed (p=0.0373;**Figure 1j**) reinstatement of extinguished seeking. All groups showed robust incubation of craving as assessed by a second cued-reinstatement test following 3months of abstinence, but responding in this test was highest in IntA animals (F_2,20_=5.703, p=0.0121;**Figure 1k**). In all tests, responding on the inactive lever did not differ significantly between groups (p’s>0.05).

Clinical reports indicate that protracted withdrawal from drug is associated with negative emotional states that drive relapse of drug taking through negative reinforcement processes (30). To test for these affective changes, after IntA/LgA/ShA treatments animals remained in their home-cage for 4wk of abstinence, and then were tested with a forced swim test (FST), saccharin preference (SPT), or open field test (OFT). Compared to ShA or LgA subjects, IntA animals exhibited heightened depressive-like behaviors on FST (time swimming: F_2,20_=4.265,p=.0305; **Figure 2a**) and SPT (F_2,21_=4.027,p=.0348;**Figure 2b**), as well as enhanced anxiety-like behavior in the OFT (F_2,18_=3.857,p=.0429;**Figure 2c**). Higher levels of anhedonia (decreased saccharin preference) were associated with higher cued cocaine-seeking following abstinence in IntA animals (R^2^=-0.914,p=.0299) but not ShA or LgA rats.

**Figure 2.**
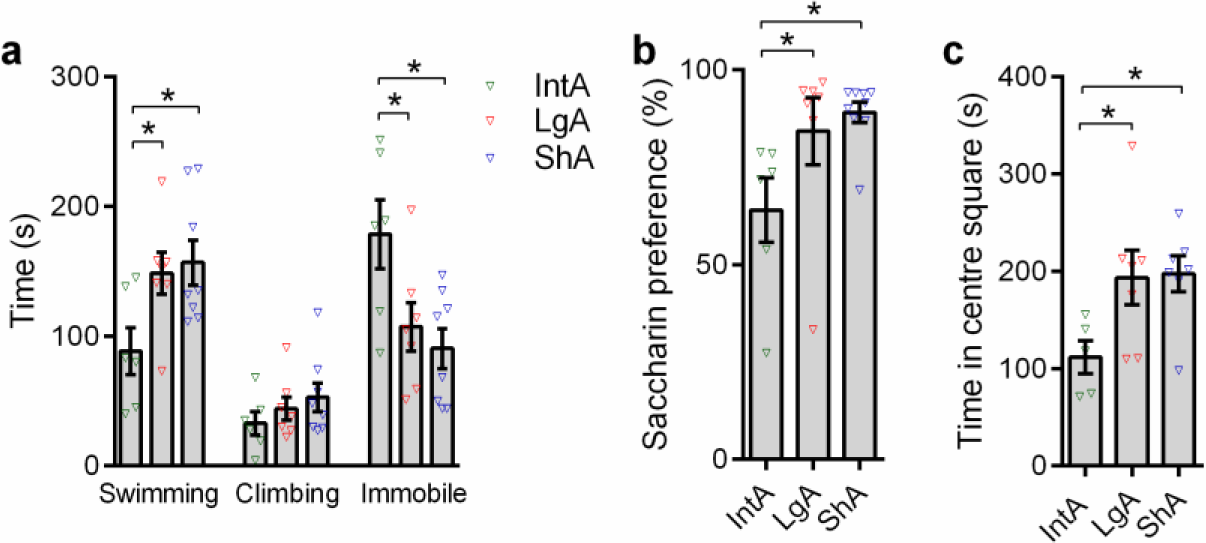
IntA is associated with the emergence of negative emotional states following 4 weeks of abstinence. (**a**) IntA animals spent less time swimming on the forced swim tests compared to LgA (t=2.348, p=.0305; Holm-Sidak post-hoc test) or ShA (t=3.132, p=.0263; Holm-Sidak post-hoc test) control animals. On the same test, IntA animals spent more time immobile compared to LgA (t=2.462, p=.0242; Holm-Sidak post-hoc test) or ShA (t=3.132, p=0.0115; Holm-Sidak post-hoc test) control rats. There was no significant difference between LgA and ShA subjects in these measures (p>0.05). IntA: n=6; LgA: n=7; ShA: n=8. (**b**) IntA animals showed lower preference for saccharin compared to LgA (t=2.121, p=0.0473; Holm- Sidak post-hoc test) or ShA (t=2.757, p=.0249; Holm-Sidak post-hoc test) rats, indicative of anhedonia. There was no significant difference between LgA and ShA subjects in this measure (p>0.05). IntA: n=6; LgA: n=7; ShA: n=7 [one ShA animal included in the FST did not complete SPT or OFT]. (**c**) IntA animals spent less time in the center square during a 15min open field test compared to LgA (t=2.412; p=.0445; Holm-Sidak post-hoc test) or ShA (t=2.525, p=.0445; Holm-Sidak post-hoc test) rats, indicating an enhanced anxiety-like state in these animals. There was no significant difference between LgA and ShA subjects in this measure (p>0.05). IntA: n=5 [one animal excluded, see *‘Statistical analyses’* section]; LgA: n=7; ShA: n=7. All analyses represent Holm-Sidak post-hoc comparisons. Error bars are s.e.m. *P<0.05.

Taken together, we conclude that the IntA paradigm is more effective than the LgA or ShA model at producing a persistent and multi-phenotype addiction-like state that reflects many of the key DSM-V diagnostic criteria for substance use disorder.

### Signaling at the orexin-1 receptor underlies the expression of an addiction-like state following IntA

The orexin-1 receptor antagonist SB-334867 (SB) reduces motivation for cocaine, particularly in animals with high motivation for drug (1, 31). Thus, here we tested effects of SB in animals that exhibit an addiction-like state following IntA. SB10 normalized β to pre-IntA values (F_2,38_=7.695,p=.0030) without affecting Q_0_ (p>.05;**Figure 3a,b**); note that this dose is 2-3X lower than those generally found to affect motivated behavior for cocaine. The behavioral effects of SB30 were also larger in IntA compared to LgA or ShA rats; SB30 had no effect on β across all LgA animals (p>.05;**Figure 3c**), but as in previous studies (25, 31), significantly increased β (decreased motivation) in a subset of high-motivation LgA rats (**Figure S2**). Neither SB10 nor SB30 significantly affected β in ShA animals (p>.05;**Figure 3d**).

**Figure 3.**
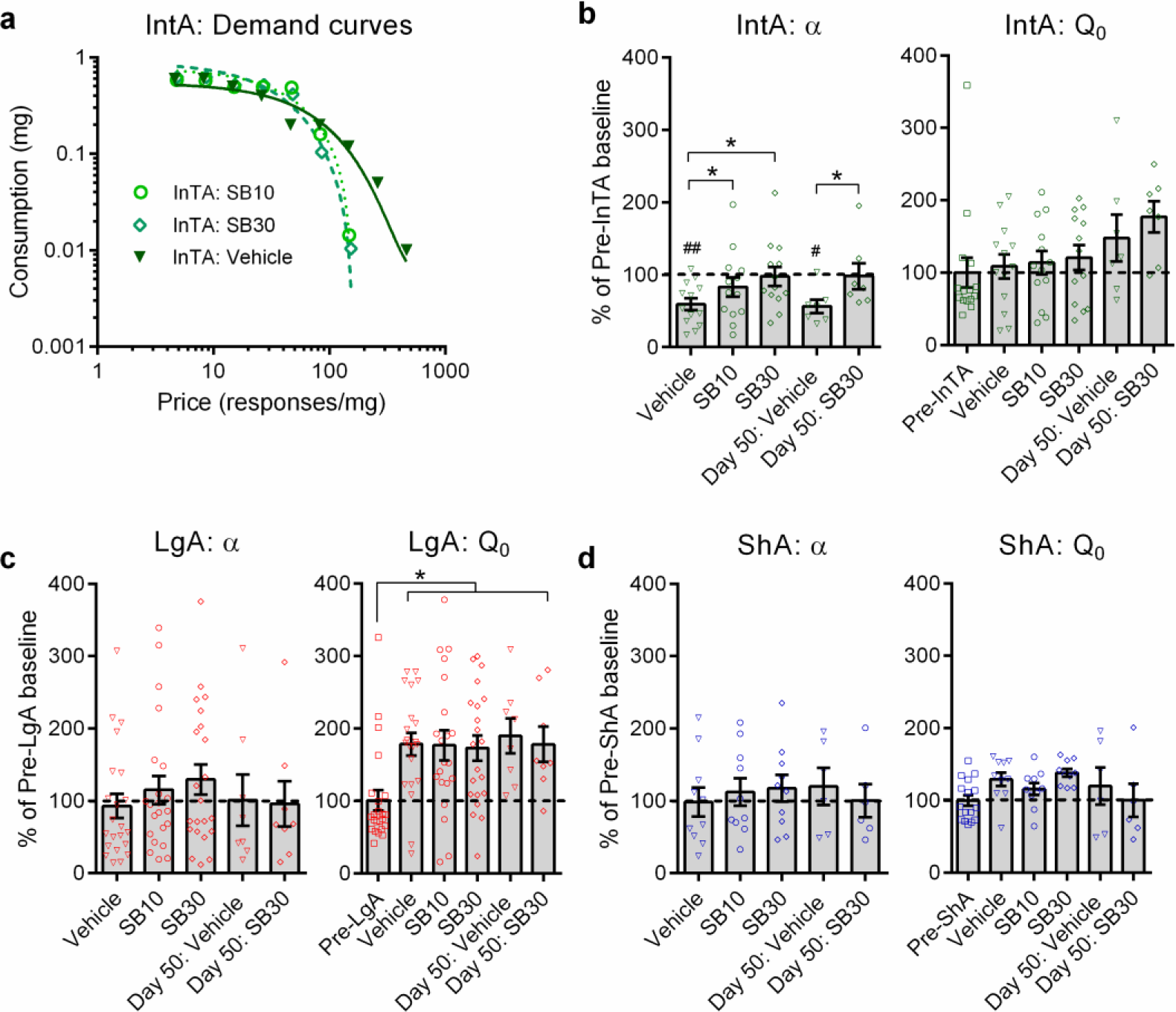
SB (OxR1 antagonist) reverses IntA-induced changes in demand (a) without affecting cocaine consumption at null effort (Q_0_). (**a**) Sample demand curves from a representative IntA animal illustrating a significant increase in demand elasticity (α; decrease in motivation) following SB30 treatment. (**b**) SB normalized β to baseline levels in IntA animals at both 10 (t=3.562, p=.0039) and 30mg/kg (t=2.545, p=.0438). SB30 remained effective at normalizing demand even when administered 50d following IntA training (W=28. p=.0156). SB had no effect on Q_0_ values in these subjects. Holm-Sidak/Wilcoxon post-hoc comparisons compared to vehicle. Veh/SB10/SB30: n=13; D50 Veh/D50SB:n=7. (**c**) SB had no effect on α or Q_0_ overall in LgA animals (p’s>0.05). Veh/SB10/SB30: n=21; D50 Veh/D50SB:n=8. (**d**) SB had no effect on α or Q_0_ overall in ShA animals (p’s>0.05). Veh/SB10/SB30: n=10; D50 Veh/D50 SB: n=6. Error bars are s.e.m. *P<0.05. #P<0.05, ##P<0.01 vs pre-IntA values.

Because the IntA-induced addiction-like state was persistent for at least 50d, we also tested the effect of SB on a at this timepoint. Remarkably, SB30 normalized a in IntA animals tested at 50d (W=28,p=.0156; **Figure 3b**), but had no effect in LgA or ShA rats (**Figure 3c,d**).

SB at both doses was also most effective on cued reinstatement behavior in IntA animals: SB10 significantly attenuated cued-reinstatement in IntA rats (F_3,35_=14.03,p=.0038), whereas this dose in ShA or LgA rats was ineffective (**Figure 4a-c**; note that previous studies indicate that 20 mg/kg is needed in ShA rats (32)). Also, SB30 more strongly suppressed cued reinstatement responding in IntA subjects than in ShA or LgA animals (IntA vs LgA or ShA;p=0.0023, Holm-Sidak post-hoc comparisons of A vehicle vs SB30). Consistent with previous studies, SB had no effect on primed reinstatement (p’s>.05;**Figure 4d-f**), indicating that the stimulus cueing properties of cocaine remained intact. SB also had no effect on non rewarded responding during reinstatement testing (**Figure S3**) or general locomotor activity (p’s>.05;**Figure 5**), indicating that SB had no non-specific effects on general activity. Together, these data indicate that rats that exhibit an addicted state following IntA are more susceptible to the anti-addiction properties of SB.

**Figure 4.**
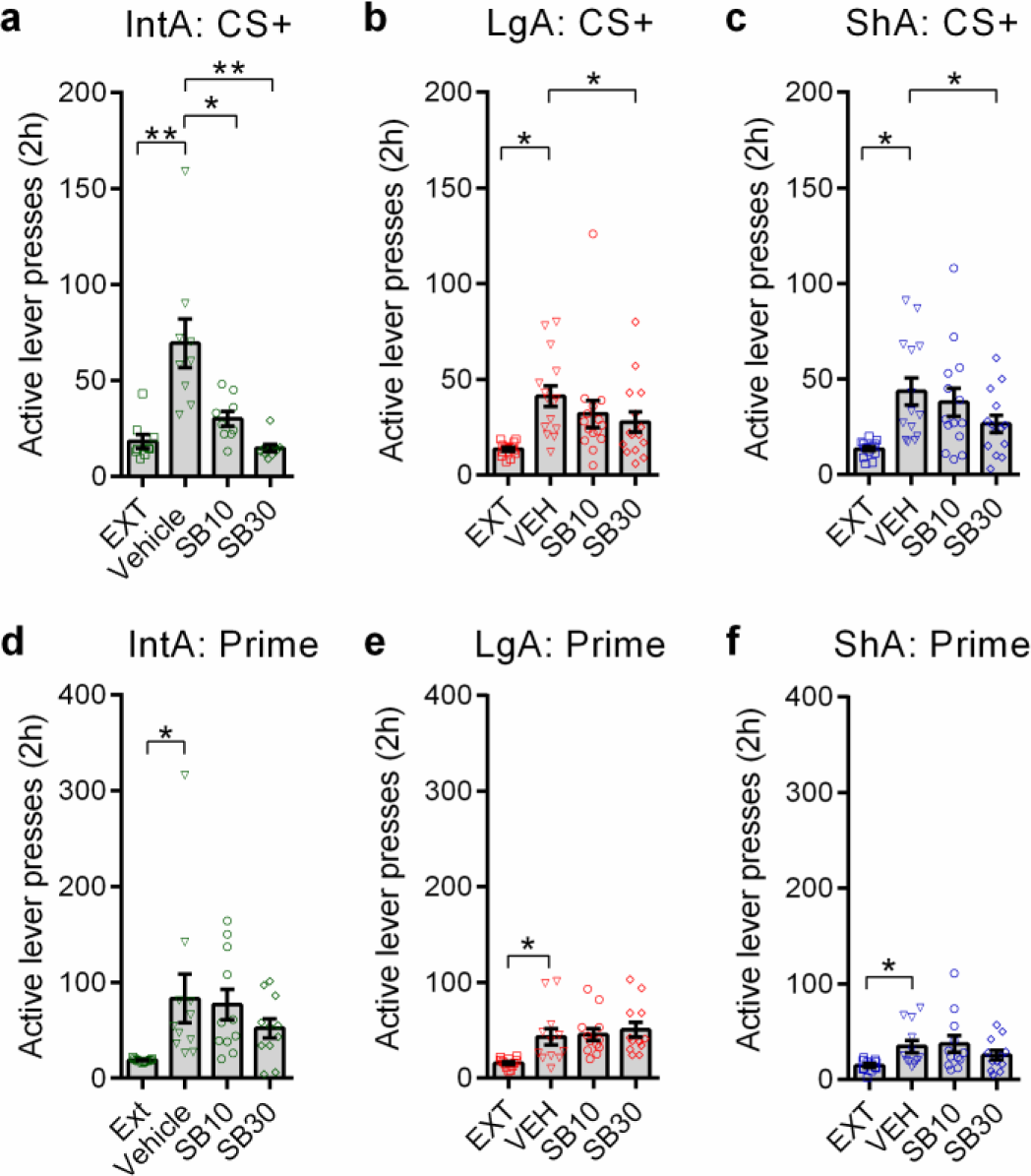
SB blocks cued reinstatement at a lower dose in IntA animals. (**a**) SB10 (t=2.750, p=.0250; Holm-Sidak post-hoc test) and SB30 (t=4.128, p=.0066; Holm-Sidak posthoc test) significantly reduced cued reinstatement of active lever responding in IntA animals. n=9. (**b-c**) Only SB30 was significantly effective at reducing reinstatement in LgA (b; t=2.147, p=.0498); Holm-Sidak post-hoc test or ShA animals (c; t=2.930, p=.0233; Holm-Sidak post-hoc test). LgA: n=15; ShA: n=14. (**d-f**) A priming dose (10mg/kg) of cocaine elicited reinstatement of active lever responding (relative to EXT levels) in IntA (a; F_3,43_=3.433, p=.0259 ANOVA; t=2.887, p=.0186, Holm-Sidak post-hoc comparison), LgA (b; F_3,47_=7.297, p=.0027, ANOVA; t=3.152, p=.0274, Holm-Sidak post-hoc comparison) and ShA rats (c; F_3,47_=4.013, p=.0291, ANOVA; t=2.887, p=.0186, Holm-Sidak post-hoc comparison. Across all groups, SB had no effect on active lever responding during these tests. IntA: n=9;LgA: n=15; ShA:n=14. Holm-Sidak post-hoc comparisons. EXT: pooled extinction values across all groups. Error bars are s.e.m. *P<0.05. **P<0.01.

**Figure 5.**
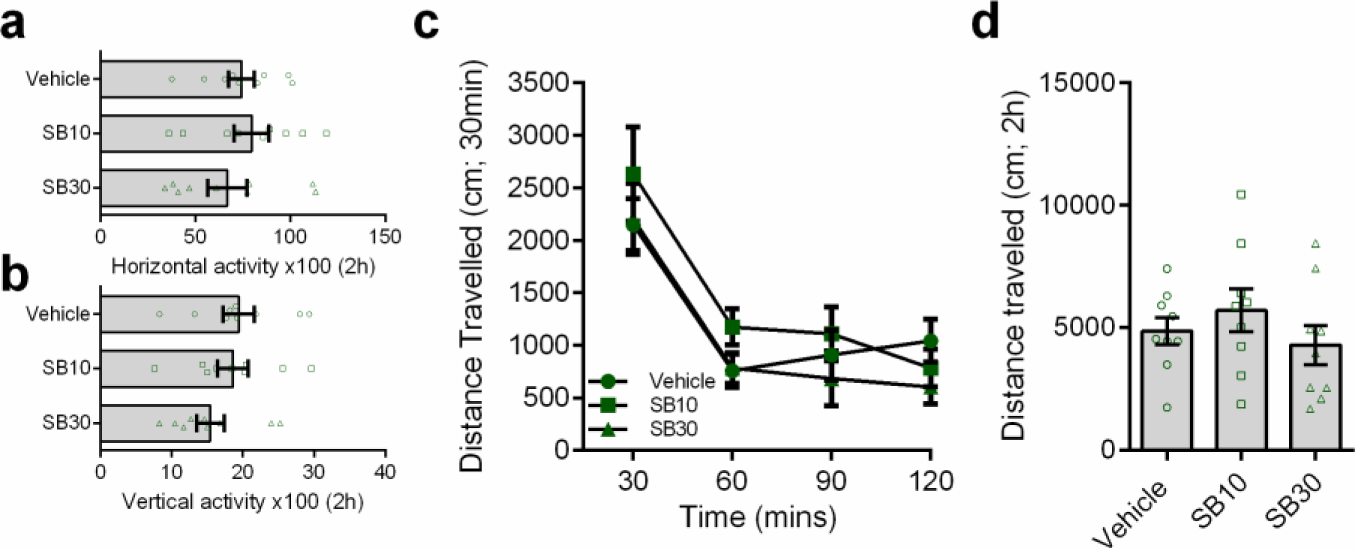
SB had no non-specific effects on general locomotor activity in animals with a history of IntA. (**a-b**)In animals with a history of IntA to coca ine, SB had no effect on horizonta l (a; F2 26=0.6678, p=.4925) or vertical (b; F2,26=1.551, p=.2450) activity over the 2h test period. n=9 (**c-d**) Similarly, SB did not affect the total distance traveled (in cm) across the 2h test period when assessed at 30min intervals (F6,72=.8465, p=.5384) or in sum (F2,26=1.315, p=.2950).n=9 N=9 for all groups. Error bars are s.e.m. *P<0.05. ****P<0.0001.

### IntA-induced addicted state is driven by a persistent augmentation of LH orexin function

To test whether the IntA-induced addicted state is associated with enhanced orexin system function, we measured orexin cell number and activity following IntA in medial (DMH/PF) and lateral (LH) orexin cell populations. Rats were sacrificed 1 or 150d following IntA training, and in both cases, 90min after being re-exposed to the cocaine self-administration environment. ShA rats were used as controls as their total daily cocaine access (1h) was identical to that of IntA animals. Immediately following IntA training (d1), we observed significantly more orexin-A immunoreactive (ir) cells in LH of IntA animals relative to ShA controls (t_14_=3.384,p=0.0045;**Figure 6a-c**). The self-administration environment also activated LH orexin neurons in IntA animals on d1 to a greater extent than ShA animals, as indicated by a larger proportion of orexin-ir neurons that co-expressed the neuronal activity marker Fos (t_14_=3.658,p=.0026;**Figure 6e**). Remarkably, the increased number (t_11_=2.570,p=.0261) and activity (t_11_=2.348,p=.0386) of LH orexin-ir neurons persisted at d150 (**Figure 6c,e**). Moreover, activity of LH orexin-ir neurons was correlated with incubation of craving as assessed by reinstatement testing following 3months of homecage abstinence (R^2^=0.870;p=0.024). Although we observed an increase in the number (t_14_=2.320,p=.0359) and activity (t_14_=3.831,p=.0018) of DMH/PF orexin-ir neurons immediately following IntA (d1) compared to ShA controls, these differences did not persist into 150d of withdrawal (p’s>.05;**Figure 6d,f**). Together, these findings indicate that the addiction-like state induced by IntA is associated with a persistent augmentation of orexin cell number and function, particularly within LH.

**Figure 6.**
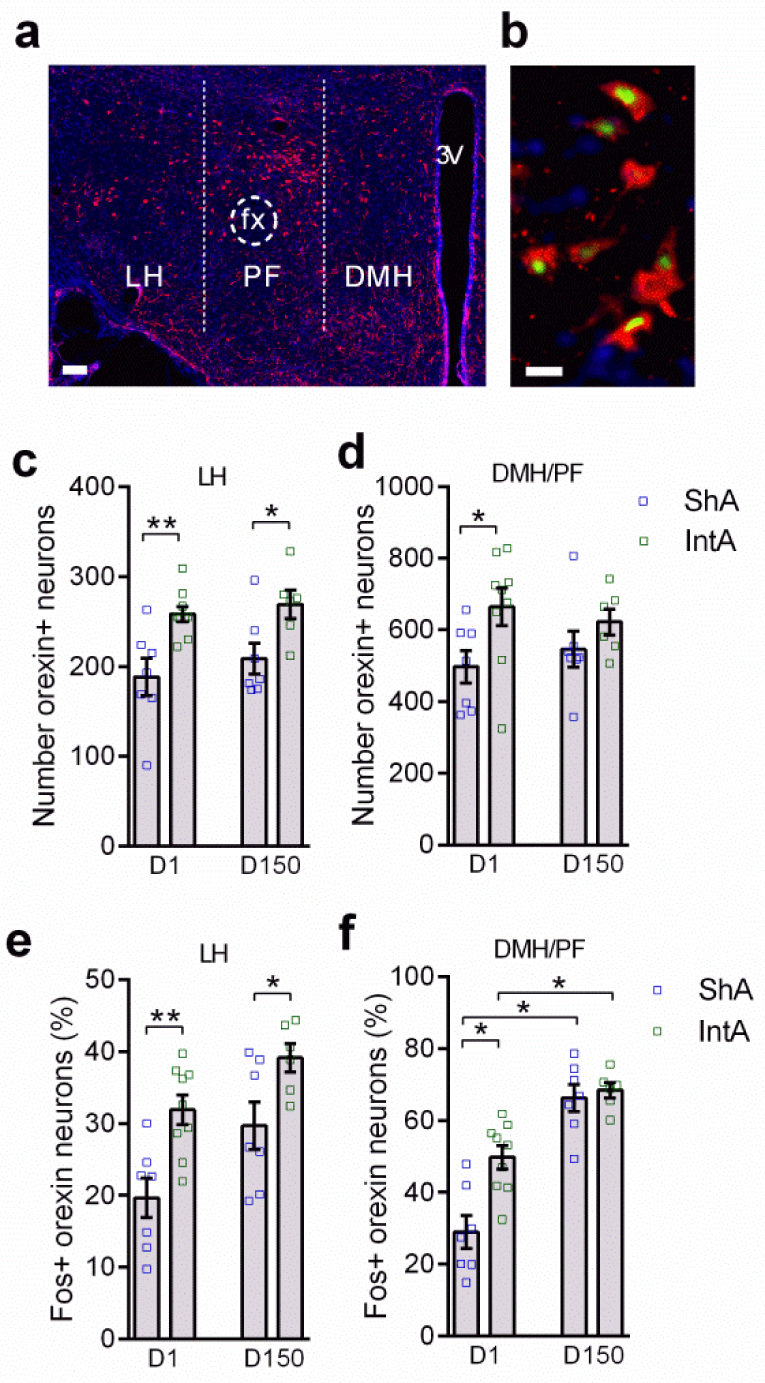
The IntA phenotype is accompanied by a persistent increase in LH orexin neuron number and activity. (**a-b**) Low (a) and high (b) magnification images of orexin-ir neurons in hypothalamus stained for orexin (red), Fos (green; Panel B only) and DAPI (blue), taken from a IntA rat. Scale bar panel A=500|am, panel B=25|am. (**c**) Compared to ShA controls, IntA animals had a significantly higher number of orexin-A-ir neurons in the LH ld (t=3.384, p=.0045; independent samples t-test) and 150d (t=2.570, p=.0261; independent samples t-test) following IntA. (**d**) IntA animals had a significantly higher number of medial (DMH/PF) orexin neurons on d1 only (t=2.320, p=.0359; independent samples t-test). (**e**) In LH, IntA animals had a significantly greater percentage of orexin-A-ir cells that were activated in response to the drug environment on days 1 (t=3.658, p=.0026; independent samples t-test) and 150 (t=2.348, p=.0386; independent samples t-test) following IntA. (**f**) In the DMH/PF, IntA animals had a significantly greater percentage of orexin-A-ir cells that were activated in response to the drug environment on day 1 only (t=3.831, p=.0018; independent samples t-test). D1: ShA, n=7, IntA, n=9; D150: ShA, n=7, IntA=6. Error bars are s.e.m. *P<0.05. **P<0.01.

### Knockdown of LH orexin cells attenuates addiction phenotype

To investigate the functional importance of augmented LH orexin signaling in the expression of an addiction-like state, we assessed the effect of knocking down orexin production in LH vs DMH/PF subregions on cocaine demand elasticity (β). We used an orexin morpholino antisense (**Figure S4a,b**), which selectively reduces orexin protein expression without affecting other neuropeptides in interdigitated neurons (8, 29). Microinjection of the morpholino antisense into the LH orexin subfield produced ~25% knockdown in LH, but not DMH/PF, orexin neurons (ti_0_=6.140,p=.0001;**Figure 7a,b**), which is roughly equivalent to the increase in cell numbers in this region following IntA. This knockdown of LH orexin was associated with a significant increase in β (decrease in motivation; t_12_=4.455,p=.0008;**Figure 7c,d**), but had no effect on Q_0_ (p>.05;**Figure S4c**). In contrast, knockdown of orexin expression in DMH/PF, but not LH, (t_8_=2.612,p=.0310) had no effect on β (p>.05;**Figure 7e**) or Q_0_ (p>.05;**Figure S4d**). Together these data show that LH, but not DMH/PF, orexin neurons are critically involved in motivation for cocaine, and indicate that the persistent changes observed in this subpopulation following IntA may underlie the expression of an addicted state.

**Figure 7.**
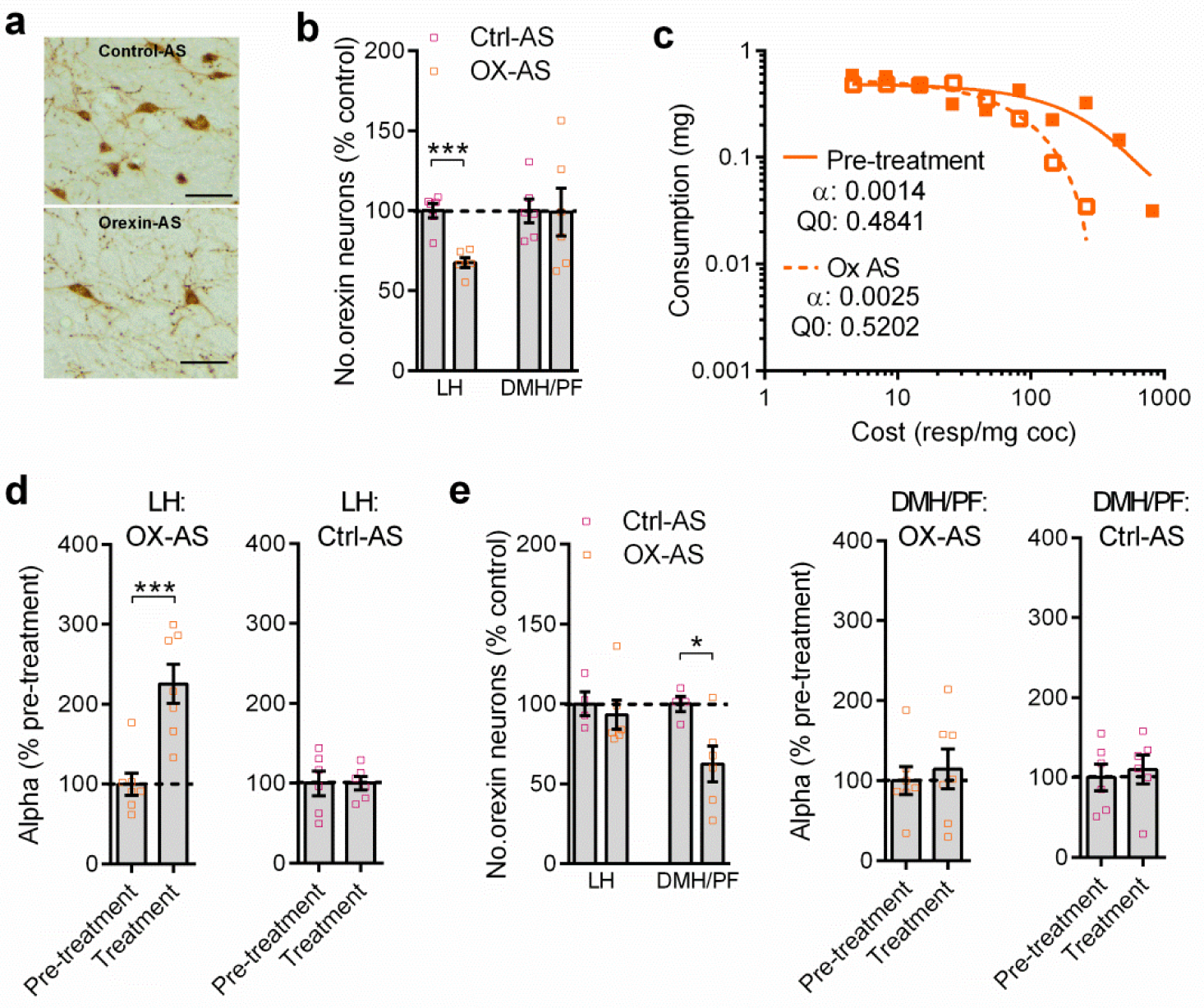
Knockdown of orexin in LH but not DMH/PF neurons attenuates addiction phenotype. (**a**) Representative photomicrographs of LH orexin-A-ir cells 6d following injection of control- (top) or orexin- (bottom) morpholino antisense. Scale bars=25μm. (**b**) Site-specific delivery of orexin-AS produced a knockdown in the number of orexin-ir neurons in LH (t=6.140, p=.0001; independent samples t-test) without affecting the number of orexin-ir neurons in DMH/PF (p>0.05). (**c**) Sample demand curves of a representative animal before (pre-treatment) and after treatment with the orexin morpholino antisense (Ox AS) in LH. (**d**) Overall, LH orexin-antisense (AS) treatment resulted in higher a values (higher demand elasticity, lower motivation; left panel; t=4.455, p=.0008; paired-samples t test). In contrast, LH control AS treatment had no effect on a values (p>0.05; right panel). (**e**) Site-specific delivery of orexin-AS produced a knockdown in the number of DMH/PF orexin neurons (t=2.612, p=.0310; independent samples t-test) without affecting LH orexin levels (p>0.05; left panel). DMH/PF Ox AS treatment had no effect on a values (p>0.05; paired samples t-test; middle panel). Similarly, DMH/PF control AS treatment did not affect a values (p>0.05; paired samples t-test; right panel). For behavioral testing of both LH and DMH/PF injections; Ox AS, n=7; Control AS, n=6. Error bars are s.e.m. *P<0.05. **P<0.01. ***P<0.001.

## Discussion

We show that IntA to cocaine promotes a state characterized by the concomitant expression of multiple addiction-relevant behaviors. This addiction-like state persists for months and is more pronounced than that induced by LgA, the current ‘gold standard’ model of addiction in laboratory animals. This IntA- induced addiction-like state is accompanied by augmented orexin system function in 3 ways: 1) increased number of orexin-expressing neurons; 2) heightened activity of orexin neurons in response to a drug- associated environment; and 3) enhanced effectiveness of an OxR1 antagonist in reducing cocaine seeking. These neuroadaptations persist long into withdrawal (a cardinal aspect of addiction) and are specific to orexin neurons in the LH orexin subfield. Moreover, we confirm that an increased number of orexin-expressing neurons in the LH orexin subfield is necessary for this pathological motivation for cocaine. These data reveal a major role for LH orexin cells in the development and expression of multiple addiction behaviors, associated with persistent plasticity within this neuropeptide circuit.

### IntA as a model of addiction in rodents

Recent work has sought to produce animal models that capture the defining features of human drug addiction (18, 33). The DSM-V defines addiction, or substance abuse disorder, as meeting 2 or more of 11 diagnostic criteria that largely focus on impaired control over drug use, including increased time and energy spent seeking and using the drug, intense craving to use the drug, an inability to reduce drug use, and social impairment and compulsive drug use including drug use despite negative consequences (34). Typically, animal studies focus on a single addiction characteristic, such as either escalation/binge patterns of drug use, enhanced motivation for drug, use despite negative consequences, or drug- seeking/relapse vulnerability. Here, we show that a composite of *all* of these endophenotypes is concomitantly enhanced *in the same animal* following IntA to a greater extent than the LgA model. Moreover, we show enhanced anxiety- and depression-like behavior following IntA, another key feature of addiction that is thought to contribute to subsequent drug use via negative reinforcement (35). Strikingly, this IntA-induced addiction-like state persists for months - enhanced motivation for drug, measured by demand elasticity on a behavioral economics task, was observed for at least 50d following IntA, and heightened reinstatement behavior was observed following 3 months abstinence. Although LgA transiently enhanced motivation for cocaine, these animals generally did not differ from ShA controls after ~1w of LgA training in terms of demand or any other measures of addiction-like behavior. This is despite a persistent (at least 50d) increase in Q_0_ among LgA animals, indicating that contrary to the prevailing view, pathological drug seeking in addiction is not necessarily a consequence of changes in the hedonic set point for cocaine following extended use (18, 19). Of note, several studies have reported an augmented addiction-like phenotype in LgA rats(19, 36–41). However, in many of these studies, behavioral testing was carried out shortly after LgA training, a time-point that may align with the transient increase in demand observed here. Together, our data demonstrate that the IntA model offers a unique behavioral paradigm to promote a robust, multifaceted addicted-like state in rats to a greater degree than the LgA model.

### Plasticity of the LH orexin system in addiction

We found that IntA produces persistent plasticity in the orexin system that is associated with the expression of multiple addiction behaviors. Specifically, we observed an increase in the number of LH orexin-expressing neurons and their responsivity to drug-associated stimuli following IntA, and an associated increase in demand for cocaine, compulsive responding for cocaine, and vulnerability to cued and drug-primed reinstatement behavior. Also, the effects of SB were greatest in IntA animals at all doses, and demand for cocaine and cued reinstatement of drug-seeking was normalized by significantly lower doses of SB than have been reported previously (25, 32, 42). Interestingly, although we previously showed that SB reduces demand for cocaine in restricted-access rats (25), we found here that SB had no effect on a in ShA animals. This may reflect greater engagement of the orexin system with longer (2hr) self-administration sessions in our previous study compared to the 1hr sessions used here.

Our results are consistent with previous findings that the orexin system undergoes substantial plasticity in response to several environmental changes. Both the number and activity of orexin neurons is higher during wakefulness than during sleep (11, 43, 44), and acute food deprivation increases levels of orexin peptide and mRNA, and the activity of orexin cells (45, 46). In addition, pre-pro orexin levels are reduced following chronic social defeat stress(46). There is also prior evidence of enduring plasticity within the orexin system following exposure to drugs of abuse, as orexin mRNA is increased in LH following chronic alcohol (16) and during withdrawal following cocaine (47). It is unclear from these reports however, whether these changes reflect neural adaptations that confer a heightened propensity to addiction. Our findings indicate that functional consequences are likely for this orexinergic plasticity. Our findings also align with previous demonstrations that LH orexin cell activity is correlated with drug-seeking behaviors (4, 48).

Plasticity in orexin system function persisted for months, consistent with the long-lasting behavioral addiction phenotype observed following IntA. IntA animals remained more sensitive to SB compared to LgA or ShA controls for at least 50d following IntA training, and the number and reactivity of LH orexin- expressing neurons remained elevated in IntA animals following home-cage abstinence (150d post-IntA).

Greater numbers of LH orexin-expressing neurons were reactive to drug-associated stimuli following prolonged withdrawal in IntA subjects, indicating that plasticity in the LH orexin system may underlie protracted relapse vulnerability following the cessation of drug use. Indeed, changes in orexin neuron function during withdrawal were associated with incubation of craving and the development of negative emotional behaviors in the same animals. Mechanisms contributing to orexin system plasticity following IntA remain to be determined. A subpopulation of LH neurons may persistently increase orexin synthesis after IntA, as previously proposed for the active phase of the diurnal cycle (43). Other evidence indicates that glutamatergic input onto orexin neurons is increased following repeated cocaine exposure (49), and this synaptic plasticity may be further enhanced by IntA. Additional work is required to examine these possibilities.

We confirm that plasticity in LH orexin neurons is critical for motivation for cocaine by showing that selective knockdown of orexin in this cell group reduces cocaine demand. This is the first functional evidence for a selective role of LH versus DMH/PF orexin neurons in drug self-administration. Together with our anatomical/activity mapping data, these results strongly support the proposed dichotomy in orexin cell function, whereby LH orexin neurons preferentially encode reward motivation, whereas DMH/PF orexin neurons process arousal and stress (1, 13). Increased numbers and reactivity of DMH/PF orexin neurons immediately after IntA training (d1) might reflect acute withdrawal-induced arousal/stress; however, our antisense results indicate that this does not contribute *per se* to the enhanced demand in these animals. In contrast, our results show that LH orexin neurons directly contribute to motivated responding for cocaine, and that enhanced function of these neurons, both immediately following IntA as well as following protracted withdrawal, contributes to the development and expression of the addicted state following IntA. This may be due to enhanced LH orexin output onto key motivational sites such as ventral tegmental area following IntA (50–54).

In sum, we show that IntA is an effective model to promote a persistently addicted-like state in rats, characterized by a comprehensive set of addiction-like behaviors that closely recapitulate key diagnostic criteria for addiction in humans. We also show that this IntA-induced addicted state depends upon enhanced orexin system function specifically within LH, such that this addicted state can be normalized via specific knockdown of LH orexin neurons or OxR1 pharmacological blockade. Thus, our data strongly implicate the orexin system in the addicted state and highlight this system as a promising target for addiction therapies.

## Acknowledgements

This work was supported by C.J. Martin Fellowships from the National Health and Medical Research Council of Australia to MHJ (No. 1072706) and HEB (No. 1128089), by a U.S. Public Health Service award from the National Institute of Drug Abuse to GAJ (R01 DA006214) and BAZ (F32 DA036995), and by the Charlotte and Murray Strongwater Endowment for Neuroscience and Brain Health (GAJ). We would like to thank Ms. Shayna O’Connor for her invaluable assistance with carrying out behavioral experiments.

